# A non-invasive method of air pollutant exposure in an avian model of development

**DOI:** 10.64898/2026.07.21.739765

**Authors:** Duncan Garner, Simon Clarke, Joshua Durrans, Prachi Stafford, Mari Herigstad

## Abstract

Air pollution is a growing public health concern. The developing fetus is particularly vulnerable, with exposure during pregnancy linked to negative developmental health outcomes. Teratogenic studies rely on the use of model organisms, such as the chick embryo, a well-established model of human development. However, existing protocols for the exposure of chick embryos to gaseous and aerosol pollutants have financial and technical limitations. Here, we present a novel, non-invasive method for the long-term exposure of chicken embryos to a gaseous toxin, carbon monoxide (CO). Exposure is performed inside airtight incubation boxes, which can be used in a standard laboratory incubator. We demonstrate reliable dosing of precise internal CO concentrations up to 200ppm, using a simple volumetric approach. Following optimization of key incubation parameters, temperature and turning frequency, we determined the impact of the system on embryo viability and development. Closed box incubation caused minor developmental delay but had no effect on chick embryo viability. Internal oxygen concentrations remained above hypoxic levels. No significant effects of exposure up to 200ppm CO were observed on embryo viability, weight or developmental stage. In conclusion, we present a non-invasive, affordable, accessible and technically straightforward exposure method for air pollutant teratogenicity studies. This method can be applied to other model systems and organisms beyond the chick embryo as well as to other gaseous and aerosol toxins. Thus, the system offers a suitable platform for future research on teratogenic doses, mechanisms and effects of air pollutants.

## Introduction

Air pollution is the leading environmental risk factor of increased mortality and reduced life expectancy in humans (Brauer et al. 2024). In 2021, over 8 million deaths and 236 million disability adjusted life years were attributable to air pollution globally (State of Global Air 2024). This places air pollution as the second leading cause of global disease burden, ahead of high systolic blood pressure, tobacco use, and dietary risks (Brauer et al. 2024). As such, air pollution is increasingly recognized as a major public health concern.

Importantly, air pollution is not a single toxin, but rather a heterogenous mix of gaseous and aerosol toxins. Common air pollutants include sulfur dioxide (SO_2_), nitrogen dioxide (NO_2_), carbon monoxide (CO), ozone (O_3_), volatile organic compounds (VOC) and particulate matter (PM) (Kampa and Castanas 2008). Extensive and ongoing research has linked these individual pollutants to negative health outcomes in humans (Huang et al. 2021; Levy 2015; Maung et al. 2022; Meo et al. 2024; Zhang et al. 2019). Despite this, our understanding of the health effects of these individual pollutants remains limited, highlighting a need for further research.

Certain groups have been identified as vulnerable to the effects of air pollutants. These include the elderly, children, pregnant women, people with co-morbidities and people of low socioeconomic status (Chen & Kan 2008; Mannucci et al. 2015; Schraufnagel et al. 2019). One group of particular interest is the developing fetus. Ambient exposure to air pollutants during gestation is associated with increased mortality, low birth weight, premature birth and a range of congenital defects (Veras et al. 2022). While teratogenic effects of individual pollutants have been identified, further research is required to better understand these effects, their underlying mechanisms and translate these findings to real-world exposures.

Ethical considerations largely prevent teratogenic studies in humans, necessitating the use of model organisms. The chicken embryo is a robust and widely used model of human development (Reyhani et al. 1980; Kirby and Waldo 1990; Männer 1993; Allen et al. 2001; Lincoln et al. 2004; Rutland et al. 2009; Goenezen et al. 2012; Wittig and Münsterberg 2020) and in the study of human disease (Chen et al. 2021; Kain et al. 2014; Ribatti 2016; Sharrow et al. 2020; Yuan et al. 2014). Chicken embryos are often favored due to their relatively low cost, short development time, large size and accessibility (Fauzia et al. 2018). Additionally, *in ovo* development of the chicken embryo removes maternal confounding effects, permitting the study of embryo-driven effects in isolation. Chicken embryo development is morphologically, molecularly and genetically similar to humans (Bednarczyk et al. 2021). Importantly, human and chick embryos share a similar program for early stages of development of key organs, such as the brain and heart (Reichert 2009).

Due to their advantages, chicken embryos have previously been used in teratogenic studies of air pollutants and toxins (Gebhardt 1968; Kotwani 1998; Matias et al. 2024). As such, multiple methods of *in ovo* exposure to air pollutants have been developed. Many studies have focused on PM as an air pollutant. Gas exchange across the chicken eggshell predominantly occurs through ‘bubble’ pores which are ∼250nm in diameter (Zhou et al. 2011), rendering the eggshell impermeable to PM. To circumvent this, PM can be suspended in liquid media and injected intra-amniotically (Binkova et al. 1999), below the embryonic disc (Simsek et al. 2012) or onto the air sac membrane (Matsumoto and Kashimoto 1986; de Siqueira Bueno et al. 2010; Jiang, Zhang et al. 2020; Shi et al. 2020). More recently, Jiang et al. (2021) developed a protocol for gaseous exposure by injecting gas into the air cell of the egg. This was primarily developed to expose chicken embryos via inhalation of the air cell environment (Jiang et al. 2020; Gong et al. 2025), which occurs late in development. While feasible, this approach is technically difficult, time-consuming and invasive, leaving the embryo susceptible to damage and infection, particularly during repeated exposure.

The most straightforward method of exposing chicken embryos to gaseous toxins is by gassing the incubation environment, often achieved using an airtight continuous flow chamber. This approach has been widely used in studies of oxygen and carbon dioxide dynamics on chicken embryonic development (Stock and Metcalfe 1984; Wineland et al. 2006; Lourens et al. 2007; Everaert et al. 2008; Molenaar et al. 2010; Nangsuay et al. 2021). Historical studies used this approach for gaseous toxins, such as CO (Baker and Tumasonis 1972; Tumasonis and Baker 1972; Alexander and Tuan 2003) and SO_2_ (Dean 1976). More recently, Dubansky et al. (2018) adapted commercial egg incubators for continuous gas flow to expose eggs to VOCs and polycyclic aromatic hydrocarbons (PAHs). However, the custom-built nature, financial costs, space and safety requirements of these continuous gas flow apparatuses make them inaccessible to many laboratories. These methods have all been limited to fewer, larger exposure chambers, preventing the researchers from studying multiple conditions or doses simultaneously. Many experimental paradigms require long-term incubation of embryos to reach the appropriate stage of development or permit chronic exposures. Dosing the incubation environment reduces the risk of embryo damage and infection, facilitating long-term and repeated exposure. However, there is a need to establish robust long-term incubation protocols to ensure reliable development within an air-tight chamber.

Here we develop a novel method for long-term exposure of chicken embryos to gaseous and aerosol toxins, to address the challenges posed by existing methods. Our method of gaseous exposure of the whole embryo is non-invasive, accessible and affordable. We develop and optimize this exposure method using CO as the gaseous toxin. CO was chosen due to being relatively understudied as an air pollutant. Additionally, historical studies have demonstrated measurable effects of whole embryo exposure to CO (Baker and Tumasonis 1972; Tumasonis and Baker 1972).

## Materials and Methods

### Chick embryo incubation

Fertilized Shaver Brown eggs (*Gallus gallus domesticus*; Medeggs Ltd, UK) were incubated in Whitley Incubation Boxes (Don Whitley Scientific, UK) at 37.5 or 38°C, custom-made to add a second valve for continuous gas flow during O_2_ monitoring. Eggs and boxes were cleaned prior to use with 70% IPA to avoid infection. Relative humidity within the incubation boxes was maintained at 80% by leaving a partially open 10ml tube of dH_2_O in the box. Eggs were placed on their side in the bottom half of an egg carton within the incubation boxes. All eggs were incubated at 17°C for 85 hours between delivery and the beginning of experimental incubation. Following experiments, all embryos were immediately terminated by decapitation. All experiments were performed under approval by the Sheffield Hallam University Research Ethics Committee (ER64386858 and ER91986798).

Incubation approaches were as follows:

1. Eggs were randomly assigned to 7-, 8-, 10- and 12-day incubation at 37.5°C with 1 turn per day, to determine attrition rate with age outside of the incubation box system.

2. Eggs were randomly assigned to 1 or 3 turns per day, and to either 37.5 or 38°C incubation temperature, to optimize incubation parameters.

3. Eggs were randomly assigned to open or closed incubation box and to 2 or 3 eggs per box, at 37.5°C with 3 turns per day, to determine effect of the incubation system on embryo development.

4. Empty incubation boxes were filled with known volumes of CO to determine if exposure concentrations could be predictably achieved.

5. Eggs were randomly assigned to CO exposure or air control using optimized incubation parameters to determine the efficacy of the protocol.

### CO exposure

Incubation Boxes were injected with CO/Air (151518-AK-C, BOC, UK) using a custom gas injecting system. First, a set amount of air was removed from the incubation box and replaced with the same volume of 1000ppm CO/Air, to achieve the desired concentration of CO within the incubation box. Embryos were exposed to CO at concentrations up to 200ppm, which is over double the highest WHO safe exposure limit of 87ppm for 15 minutes (World Health Organization 2021) and much higher than exposures showing significant effects on the embryo (Matias et al, 2024) and thus expected to elicit an effect. The gas within the incubation box was mixed using the syringe to pump a volume of at least 250ml in and out of the incubation box 3 times. During CO exposure, eggs were turned 180° three times per day at 9:00, 13:00 and 17:00, with incubation boxes opened for egg turning and immediately resealed and refilled with CO.

### Internal CO measurement

CO concentrations were measured using EasyLog CO data loggers (EL-CO-USB, Lascar Electronics, UK) and 48*i*-TLE Enhanced Trace Level CO Analyzer (48I-TLE, Thermo Scientific, UK). CO data loggers were placed in closed incubation boxes and CO injection was performed to achieve target CO concentration. After 5 minutes, boxes were opened and CO concentrations returned to baseline (0ppm) for 5 minutes. This process was repeated sequentially for increasing target CO concentrations. TLE analysis was performed according to the manufacturer’s protocols, adapted for a closed loop system. Briefly, AtmoGen Jar Connection Tubing (A10155, Don Whitley Scientific, UK) was attached to the sample and exhaust ports to establish a closed loop system, with valved ports for introduction of incubation boxes into the loop. The system was preloaded with CO/Air at the target CO concentration, to flush air from the system. The incubation box was then introduced to the closed loop, and CO concentration was recorded once the reading stabilized. The Gas Analyzer was calibrated with 180ppm CO/Air before each experiment, according to the manufacturer’s protocol.

### Internal oxygen measurement

Oxygen concentrations were measured using the Kane 975 Industrial Flue Gas Analyzer (KANE975, Kane International, UK, kindly provided as a gift by Kane International Ltd) adapted by attaching AtmoGen Jar Connection Tubing (A10155, Don Whitley Scientific, UK) to the inlet and outlet, to enable compatibility with ports of incubation boxes. After 16 hours of incubation, incubation boxes were removed from 37.5°C and placed at room temperature. The instrument was allowed to equilibrate with ambient air for 2 minutes. Following equilibration, the instrument inlet and outlet were attached to the valves of the incubation box to form a closed loop system. O_2_ concentration was recorded as the lowest reading measured by the gas analyzer. The gas analyzer was allowed to re-equilibrate with ambient air back to baseline readings between measurements.

### Chick embryo measurement

Chick embryos were removed from eggs by the following procedure. Briefly, a small hole was made in the egg using a dissecting needle (12030, Bochem, Germany). Viable embryos were identified through observed heartbeat or motility *in ovo,* following windowing. Dead embryos were staged (see below) and marked as non-viable. Embryos that had died at a very early stage of development were identified through visible remnants of blood circulation development. Using forceps, the hole was widened by removing shell until the embryo could be removed using a spoon. The embryo was placed in ice cold PBS and extraembryonic membranes removed using forceps. The embryo was then moved to a dry glass petri dish and weighed using an analytical balance (PX224/E, OHAUS, Switzerland), before being returned to PBS and staged. Staging was performed according to Hamburger Hamilton (HH) staging system for chick embryonic development (Hamburger and Hamilton 1951).

### Statistics

Graphs were generated and statistical analysis were performed using GraphPad Prism 10 software (GraphPad Software, USA). Normality was assessed using the Shapiro-Wilk test. Ordinal and non-parametric data was analyzed using the Kruskal-Wallis test with Dunn’s multiple comparisons test. Parametric data was analyzed using one-way ANOVA with Tukey’s or Dunnett’s multiple comparisons test, for incubation and CO exposure experiments respectively. For statistical analysis of CO exposure experiments, all conditions were compared to the 0ppm condition.

## Results

The percentage of fertile eggs across all experimental batches (N=133) was 91.0%+/-8.7, ranging from 75% to 100% fertile eggs per batch. Only fertile eggs were included in the further analysis.

### Incubation of chicken embryos

Initially, chicken embryos were incubated at 37.5°C with 1 turn per day. Under these conditions, viability was reduced with increasing age, with the largest drop seen from day 7 (75.0% viable, N=12) to day 8 (69.2% viable, N=13; Fig. 1). There was no difference in viability between D10 (66.7% viable, N=9) and D12 (66.7% viable, N=9).

**Figure 1.**
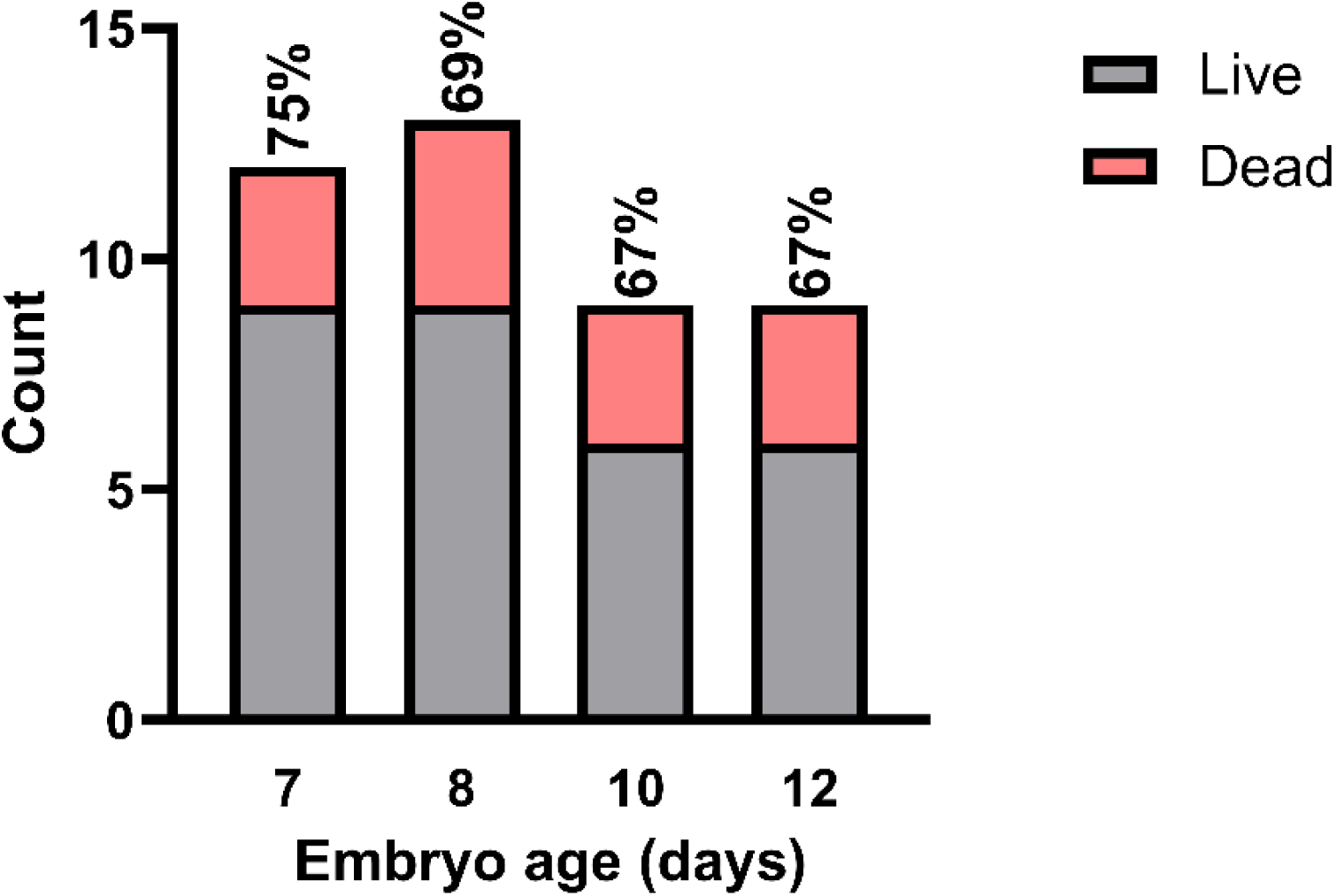
Chicken embryo viability decreased up to 10 days of incubation. Viability of chicken embryos incubated at 37.5°C with 1 turn per day. Percent viability denoted for each developmental day.

Subsequent experiments used embryos aged D10, capitalizing on the observed preserved viability following this age while also keeping to a lower stage of gestation to compensate for any potential accelerative effects of CO treatment on development.

The effect of temperature and number of turns per day on viability and developmental stage is shown in Table 1.

**Table 1.** Viability and developmental stage of chicken embryos incubated at different temperatures and turning protocols. Biological replicates (N) denoted for each group.

| Temperature (°C) | # turns / day | N | Viability (%) | HH35 (%) | HH36 (%) |
| --- | --- | --- | --- | --- | --- |
| 37.5 | 1 | 13 | 76.9 | 70.0 | 30.0 |
|  | 3 | 14 | 100.0 | 28.6 | 71.4 |
| 38.0 | 1 | 13 | 84.6 | 0.0 | 100.0 |
|  | 3 | 12 | 91.7 | 0.0 | 100.0 |

Turning affected viability, with three turns per day showing greater embryo survival than one turn per day across temperatures. The greatest effect was seen at 37.5°C, where turning thrice gave a rise in viability of 30.0% compared to turning once, whereas this rise was 8.3% for 38°C. Conversely, there was no apparent effect of temperature on viability. Turning once at 37.5°C had the lowest viability and lowest developmental stage of all conditions. Non-viable embryos were staged at HH27.3 +/-1.0 (range 26 - 28) across the eggs turned once, and at HH30 for eggs turned thrice. Linear regression analysis showed that turning frequency was a significant predictor of HH stage (p=0.04). Neither turning frequency nor temperature were predictors of viability. Subsequent incubations below used 37.5°C and turning thrice per day to maximize viability as well as developmental stage.

### Incubation of embryos in incubation boxes

During air exposure, overall viability did not differ between eggs incubated in closed incubation boxes (92% viability, N=12) and open incubation boxes (92% viability, N=14).

Optimization of number of eggs per incubation box was kept to (open box) control, 2 eggs per (closed) box and 3 eggs per (closed) box. 2 and 3 eggs per box were chosen as 1 egg per box would be prohibitively low throughput for an experimental setting and 4 or more eggs per box resulted in severe hypoxia, reduced viability and major developmental delay (preliminary observations, data not shown). Both 2 and 3 egg boxes had higher viability (100%) than open box controls (88%, Fig. 2A). 86% of open box controls were found to be HH36, an expected Hamburger-Hamilton stage for D10 of incubation at 37.5°C (Table 1). Both 2 and 3 egg closed box conditions were delayed compared to open box control (29% at HH36, p = 0.006 and 33% at HH36, p=0.025, respectively, Fig. 2B). This was reflected in embryo weight (Fig. 2C). There was no significant difference between 2 and 3 egg closed boxes for neither stage nor weight.

**Figure 2.**
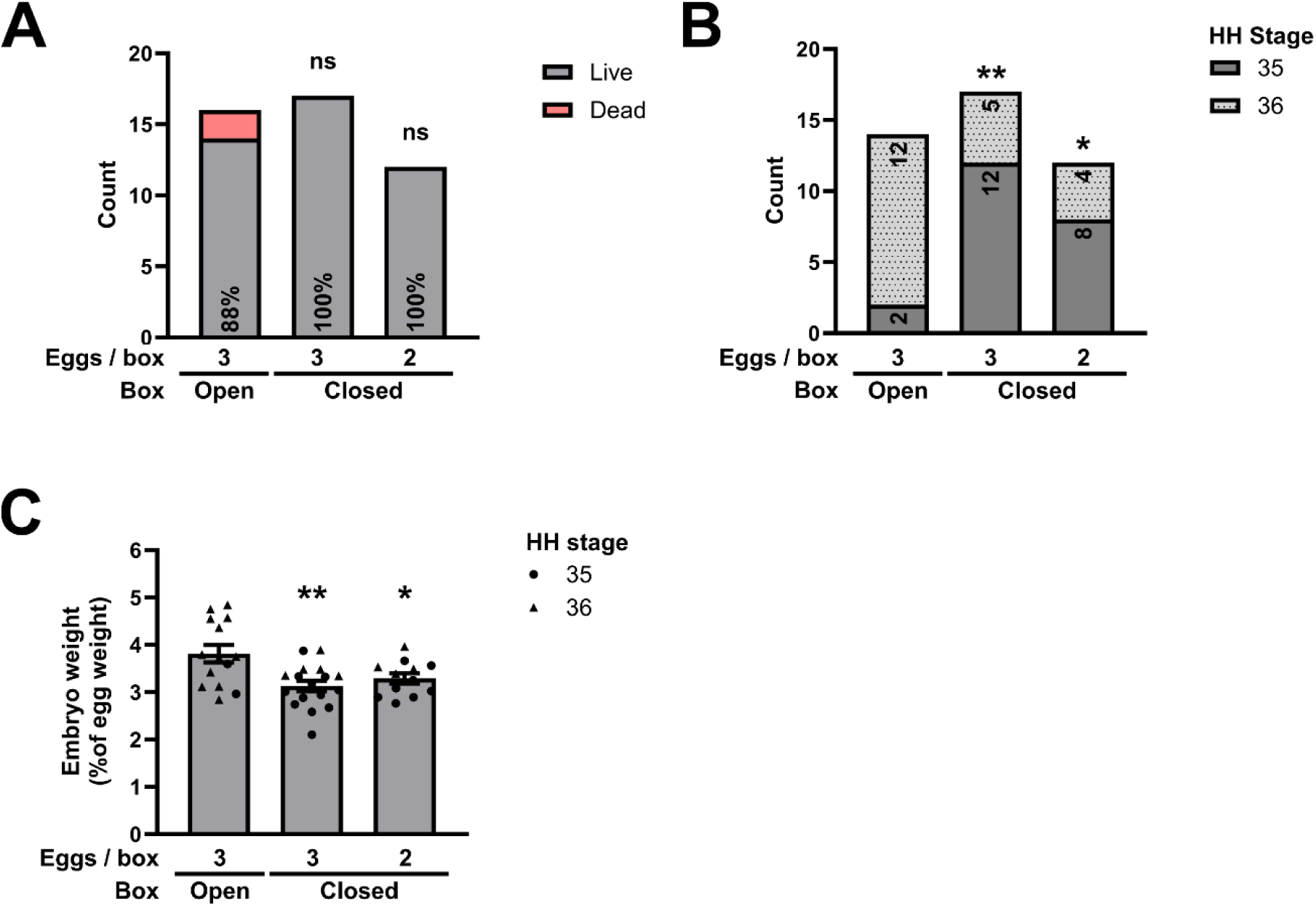
Closed box incubation causes minor developmental delay. A) Viability of D10 chicken embryos incubated in open box or closed box containing 3 or 2 eggs. Percentage viability denoted for each condition. Statistical testing is Kruskal-Wallis test with Dunn’s multiple comparisons test. B) Hamburger-Hamilton stages of D10 chicken embryos incubated in open boxes or closed boxes containing 3 or 2 eggs. Statistical testing is Kruskal-Wallis test with Dunn’s multiple comparisons test. *=p<0.05, **=p <0.01. C) D10 chicken embryo weight as a percentage of whole egg weight. Error bars represent SEM, shape represents Hamburger-Hamilton stage. Statistical testing is one-way ANOVA with Tukey’s multiple comparisons test. *=p<0.05, **=p<0.01. N=3.

Hypoxia was assessed by measuring the concentration of oxygen within the closed box environment at the end of the overnight closed incubation (the lowest expected O_2_ concentration during experimental protocols, Fig. 3).

**Figure 3.**
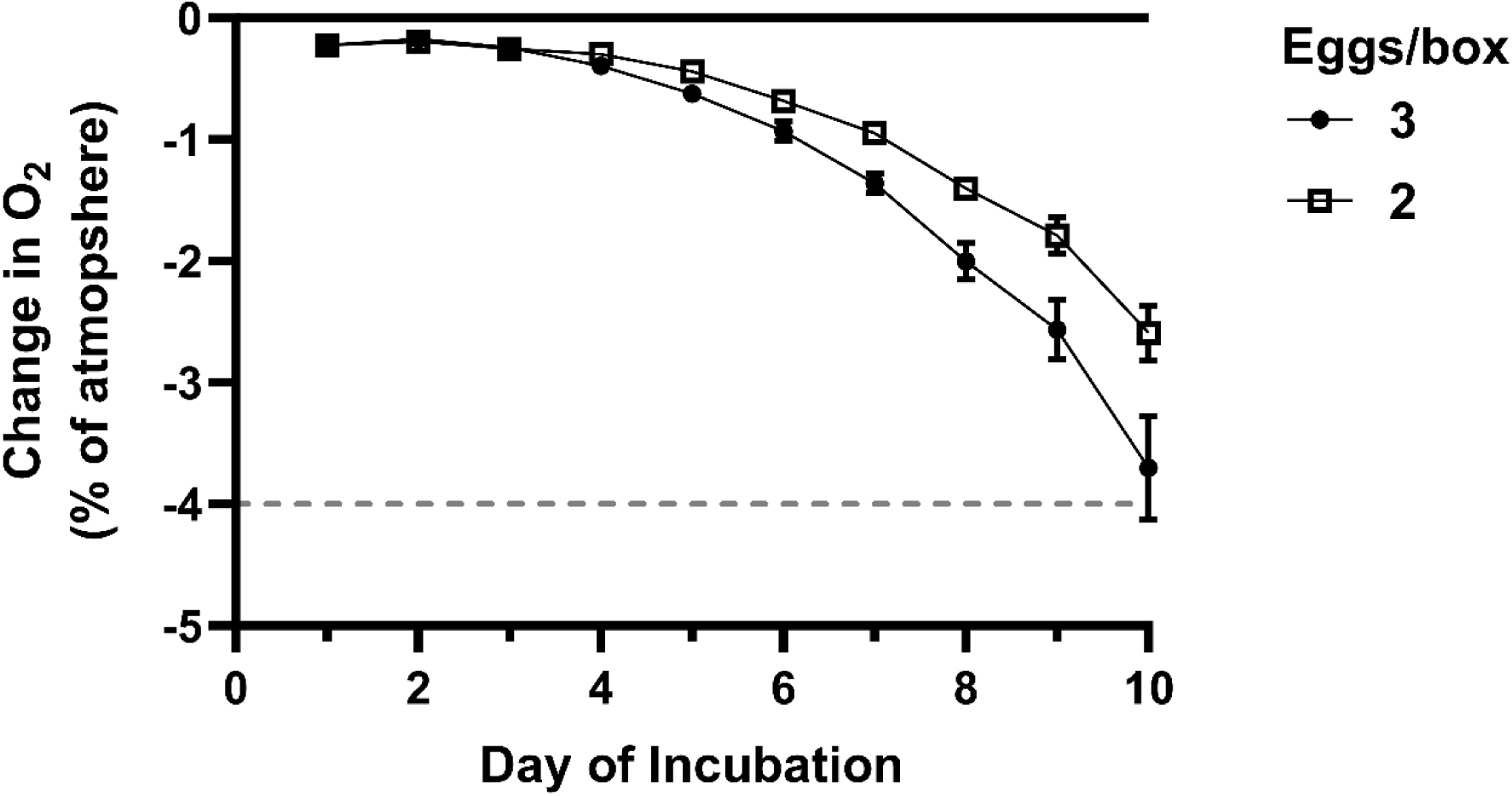
Closed overnight incubation of 2 or 3 eggs leads to non-toxic hypoxia. Change in O2 concentration (% of atmosphere) after 16 hours of continuous closed incubation. Data shown for every day of 10-day incubation. Points represent mean, error bars represent SEM, dashed line represents 17% O2. N = 3.

Past D2, atmospheric oxygen decreased exponentially with day of incubation, with boxes containing 2 or 3 eggs diverging at D4 of incubation. By D10 of incubation, O_2_ had been reduced by 2.59 and 3.70% of atmosphere in boxes containing 2 and 3 eggs, respectively. Atmospheric oxygen for both conditions was above 17% O_2_.

### Validation of CO injection protocol

The concentrations of CO tested were 0ppm (air control), 10ppm, 20ppm, 40ppm, 80ppm, 120ppm and 200ppm (Fig. 4). Volume of CO injected was calculated by first determining the actual volume of air in the box, equaling the volume in the incubation box plus the volume in the connecting tube, minus the volume of the experimental content (i.e. the combined volume of the eggs). The volume of CO mix needed to reach target concentrations can be calculated by dividing the target concentration by the concentration of CO in the experimental gas mix and multiplying this by the actual volume in the box. This method reliably produces the target concentrations required, and these remained stable following a brief overshoot in CO prior to gas mixing taking place within the box (Fig. 4A, B).

**Figure 4.**
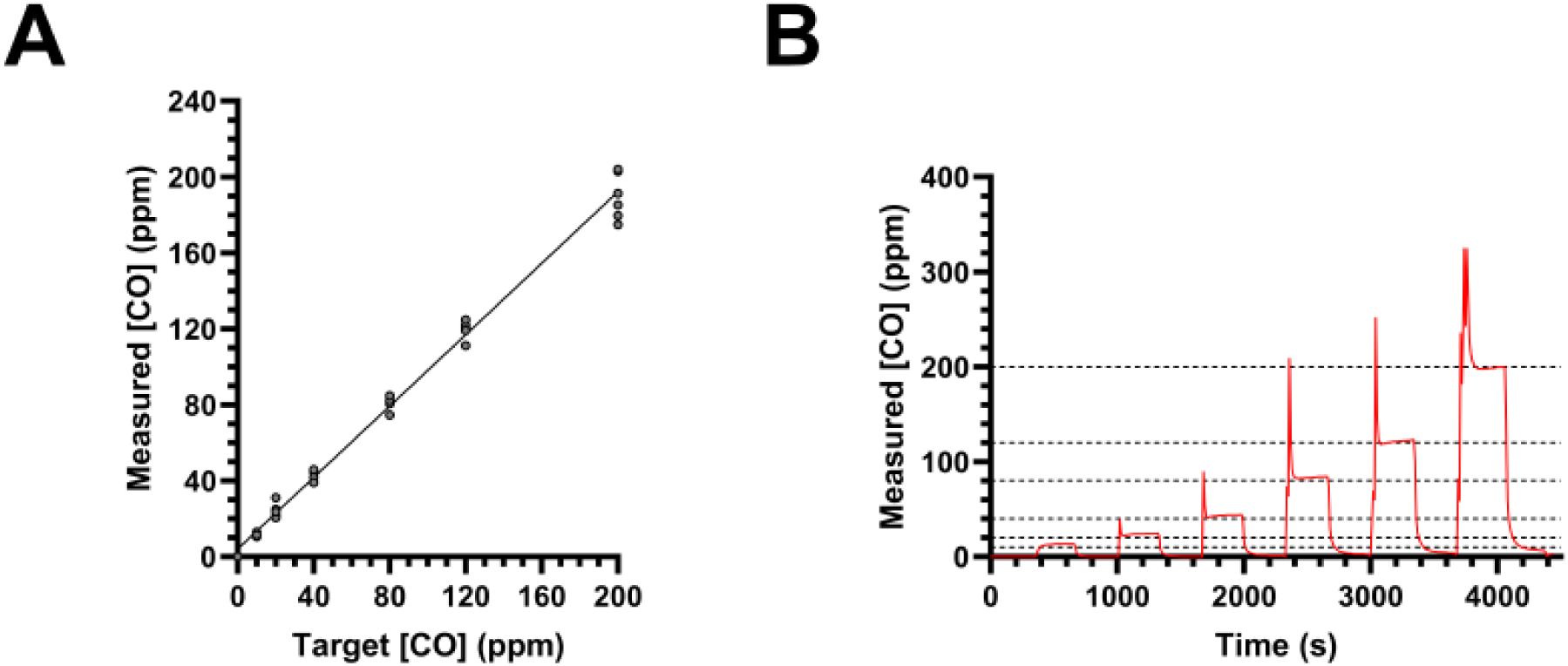
CO injection protocol achieves target CO concentrations in incubation boxes. A) Internal CO concentration measured, after injection, at each target CO concentration using 48i-TLE Enhanced Trace Level CO Analyzer. N = 6 B) Internal CO concentration (ppm) of incubation box during 5-minute exposure at each target concentration (dashed lines). Each exposure sees an initial overshoot prior to gas mixing fully with the air in the box. Data plotted as mean reading from 3 loggers.

### Chronic exposure of chicken embryos to CO

The concentrations of CO used were 0ppm (air control), 10ppm, 20ppm, 40ppm, 80ppm, 120ppm and 200ppm. All CO-containing boxes were compared against air control, which was also compared to an open box incubated under identical conditions.

Embryo viability, developmental stage and weight are shown in Figure 5. Embryos in closed boxes without CO had 100% viability. Embryo viability was reduced at all concentrations of CO, with 20 and 40ppm showing the lowest viability at 86% and 85%, respectively. Viability in CO-containing boxes ranged 85% – 93% and was comparable to open box control (88%). Significantly delayed HH developmental stage (p<0.0001) and decreases in embryo weight (p<0.0001) were observed in embryos incubated in all closed boxes compared to open box controls. Neither HH stage nor embryonic weight was significantly affected by exposure to CO at any concentration (p=0.99 and p=0.96, respectively).

**Figure 5.**
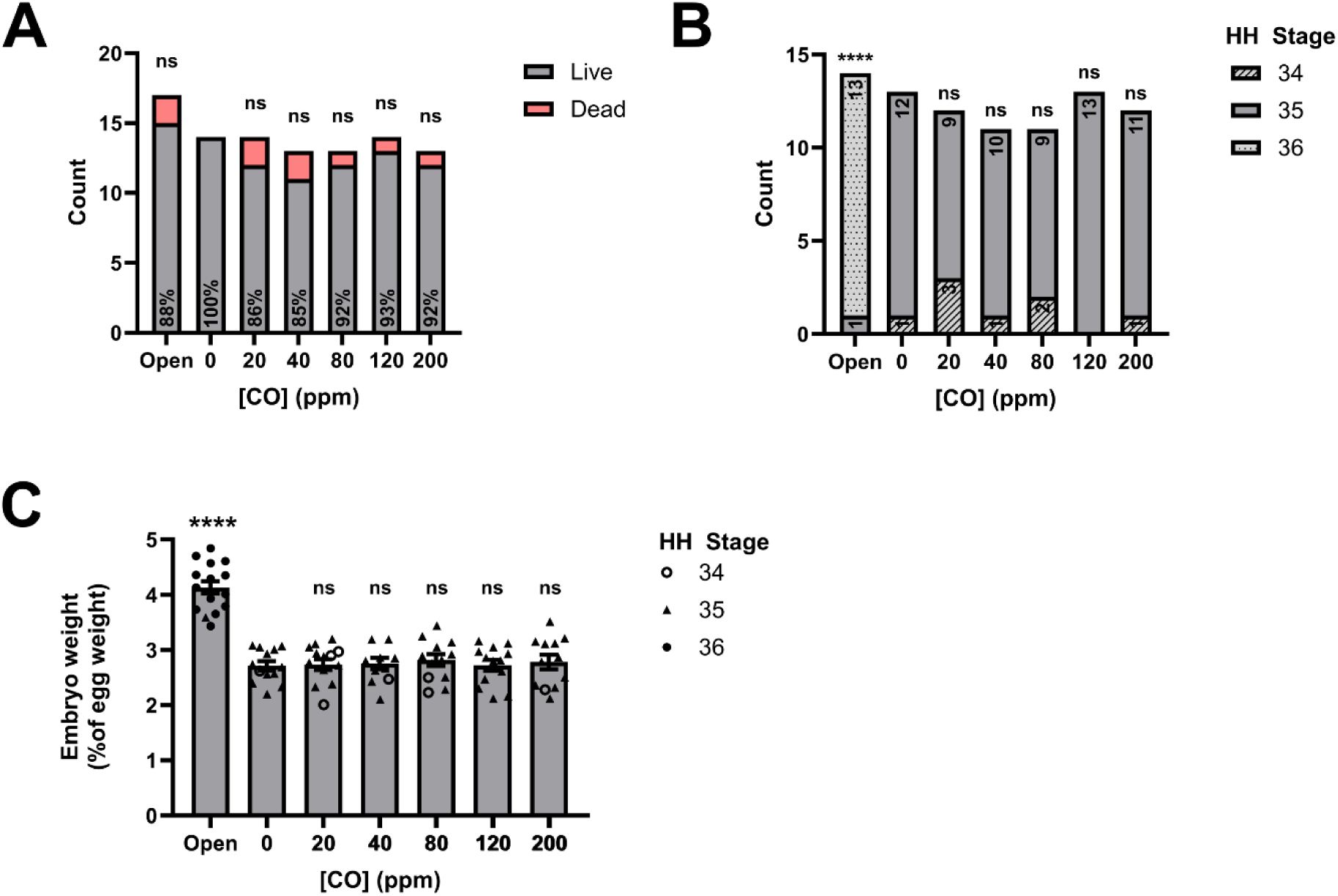
Chronic CO exposure does not have adverse effects on chicken embryos. A) Viability of D10 chicken embryos exposed to up to 200ppm CO in closed boxes. Percentage viability denoted for each condition. Statistical testing is Kruskal-Wallis test with Dunn’s multiple comparisons test of all conditions to 0ppm. B) Hamburger-Hamilton stage of D10 chicken embryos exposed to up to 200ppm CO in closed boxes. Statistical testing is Kruskal-Wallis test with Dunn’s multiple comparisons test of all conditions to 0ppm. ****=p<0.0001. C) D10 chicken embryo weight as a percentage of whole egg weight. Error bars represent SEM, shape represents Hamburger-Hamilton stage. Statistical testing is one-way ANOVA with Dunnett’s multiple comparisons test of all conditions to 0ppm. ****=p< 0.0001. N=5.

O_2_ concentration within the closed box environment was measured at the end of the overnight closed incubation. O_2_ concentration decreased exponentially past D2 of incubation for all conditions, but no significant differences were observed between air control and exposure to CO at any concentration, at any day of incubation (Mixed-effects analysis with Dunnett’s multiple comparisons test, p<0.05, Fig 6.).

**Figure 6.**
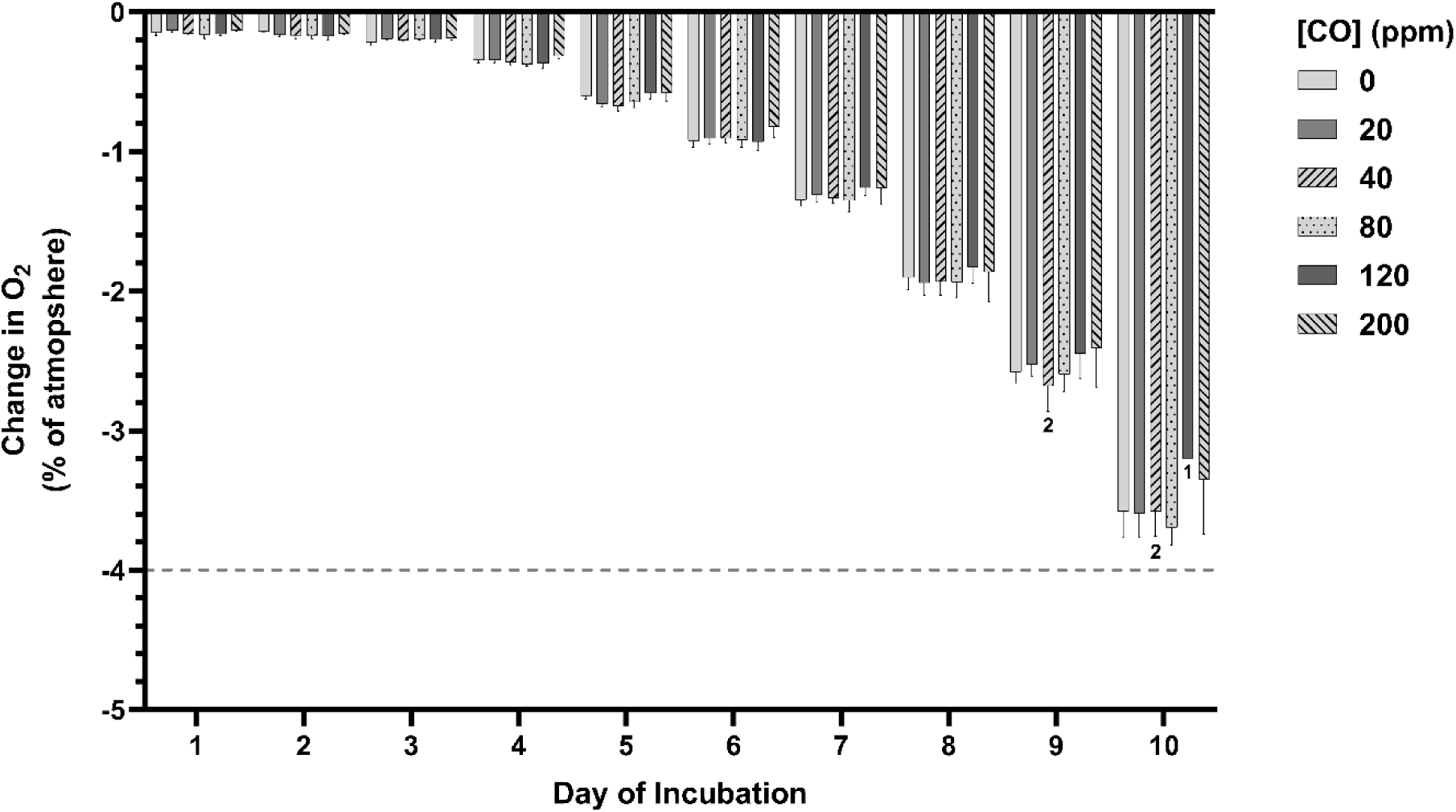
CO Exposure does not significantly change severity of hypoxia from overnight closed incubation. Change in O2 concentration (% of atmosphere) after 16 hours of continuous closed incubation. Data shown for every day of 10-day incubation in up to 200ppm CO. Bars represent mean, error bars represent SEM, dashed line represents 17% O2. Bar labels represent N where N < 3 due to reduced fertility and/or viability.

## Discussion

This paper outlines a novel method of gaseous exposure in a widely used model of human development, which overcomes many of the issues posed by commonly used methods. Using compact, sealed boxes compatible with standard cell culture incubators, commonly found in research laboratories, this method is accessible to most research laboratories. Furthermore, this method reduces the exposure volume, reducing impact or risk to the laboratory environment and researchers.

Here we outline a feasible protocol for incubating chick eggs in airtight incubation chambers. Importantly, we observe that viability is higher in eggs turned more frequently, and that the stage of the embryo can be predicted by the frequency of egg turning. Turning eggs during incubation is essential for normal embryo development. Hens turn their eggs up to 1.5 times per hour, and artificial incubation without regular turning has been associated with poor embryonic survival (Deeming 2009). This is likely due to key anatomical and physiological changes, such as reduced sub-embryonic fluid and delayed development of the area vasculosa, which are dependent on egg turning (Deeming 1989; Deeming 2009). Reduced turning is also associated with late mortality (Deeming 2009), as seen in the current study where mortality stage of non-viable eggs turned once was HH26-28. Here, viability was consistently above 90% for all control experiments where eggs were turned thrice. Importantly, our optimized incubation protocol confers increased viability compared to previous studies using airtight incubation chambers (Sharma et al. 2006), to a similar level as studies using hourly turning protocols (Gustafsson and Jacobsson 2019; Adriaensen et al. 2022).

This work provides a validated protocol for exposure of chicken embryos in teratogenic studies. Chicken embryos are an established model in teratogenicity studies (Jelinek 1982; Wachholz et al. 2021), but their use in studies of gaseous and aerosol teratogens has been hampered by technical limitations of exposure protocols. However, mounting evidence of negative developmental outcomes of human exposure to air pollutants and toxins (Veras et al. 2022) highlights the need for further study of such teratogens. We provide a straightforward and validated protocol for exposure of chicken embryos to gases and aerosols, to facilitate such studies.

Outside of teratogenicity studies, chicken embryos are also widely used in human health research. Chicken embryos are widely used in cardiovascular (Katrancioglu et al. 2012; Mangir et al. 2019) and cancer (Li et al. 2015; Mitrevska et al. 2023) research, using chorioallatoic membrane assays to study angiogenesis, tumor formation and metastasis. The chicken embryo is also an established model for ocular disease research, due to its large eyes, ease of handling and relative similarity to humans (Wisely et al. 2017). Given the existing evidence of air pollution-mediated effects on cardiovascular health (Bourdrel et al. 2017), cancer risk (M. Wang et al. 2025), and in several ocular diseases (Grant et al. 2022; Z. Wang et al. 2025), our exposure method may provide research opportunities in these fields. The chicken embryo is also emerging as a model of ischemic stroke (Fauzia et al. 2018) and the microbiota-gut-brain axis (Huang et al. 2022), highlighting its developing potential in human health research.

The method also offers scope for use with other small model organisms, such as *Caenorhabditis elegans* and *Drosophila melanogaster*. *Drosophila* are a well-established model organism, largely used for their genetic tractability, that have already been used to study the biological effects of air pollutants (Eom et al. 2017; de Santana et al. 2018). Furthermore, *Drosophila* are particularly well used to study aging and neurodegeneration (Prüßing et al. 2013; Aryal and Lee 2019; Bolus et al. 2020; Sharpe et al. 2021), for which there is mounting evidence of an accelerative effect of air pollution (Jankowska-Kieltyka et al. 2021). Using our method, large numbers of flies could easily and simultaneously be exposed to air pollutants. Such a system could provide a reliable medium-throughput tool for dosing and screening experiments.

The system is suitable for exposure of cultured mammalian cells and organ slices, which would facilitate the study of cell- and tissue-specific effects of gaseous and aerosol toxins. These models would be invaluable in attempting to understand the biological mechanisms underlying the human health effects of air pollutants. With established cell and organ slice models for a wide range of human cells, tissues and organs, this system could be used to study the effects of toxins throughout the human body. Cultures are typically carried out in liquid medium, presenting a challenge for aerosol and gaseous exposures. Exposure is often achieved by suspension of toxic particles in the culture medium (Fujii et al. 2001; Montiel-Dávalos et al. 2010; Gillespie et al. 2013) or by incubation in a dosed environment (Poma et al. 2017; Dreyer- Andersen et al. 2018), which can be easily performed with our exposure method. To accelerate the dosing of the medium, gas can be bubbled through the medium before incubation (D’Amico et al. 2006). With our method, pre-equilibration of media can be achieved with precision by incubating media in dosed boxes.

Importantly, we have demonstrated that continuous exposure of chicken embryos to up to 200ppm CO does not significantly affect viability, developmental stage or embryonic weight at D10 of incubation. To our knowledge, only 2 previous studies have examined the effects of CO exposure on chicken embryonic health and development (Baker and Tumasonis 1972; Matias et al. 2024). Consistent with our findings, Baker & Tumasonis (1972) observed no effect of continuous exposure to 100ppm CO on hatchability, another measurement of viability. Exposure to 425ppm CO was found to decrease hatchability and significantly increase embryonic weight at days 10 – 16 of incubation. Matias et al. (2024) identified no significant changes in embryonic weight and developmental stage in D9 chicken embryos continuously exposed to up to 18ppm CO, although noted a trend towards reduced weight and demonstrated significant impact on cardiac wall structure. Similar, tissue-specific developmental effects of maternal low-dose CO exposure have been reported in humans (Dadvand et al., 2011; Dix-Cooper et al., 2012). This suggests that high doses of CO are required to alter gross morphology in the chick embryo, but that detrimental changes may occur at the tissue and cellular level at much lower doses.

The above observations are supported by studies on effects of CO exposure on birth weight in both human epidemiological studies and *in vivo* animal studies. While there is some discrepancy between studies, there has been a consistently identified association of maternal CO exposure with an increased risk of low birth weight in humans (Ritz and Yu 1999; Maisonet et al. 2001; Salam et al. 2005; Morello-Frosch et al. 2010; da Silva et al. 2014; Giovannini et al. 2020). Similarly, direct exposure studies in rodent models have also linked CO exposure to birthweight. Exposure to 150ppm reduced birthweight and postnatal weight gain in rats (Fechter and Annau 1977; Mactutus and Fechter 1984), while lower exposures up to 90ppm did not affect birthweight (Garvey and Longo 1978). Exposure to ≥90ppm CO reduced birthweight in rabbits (Astrup et al. 1972), while in mice, birthweight was reduced by exposures >300ppm (Venditti et al. 2011). It is difficult, given the variation in concentration and duration of exposure, to determine why some studies observed weight changes with lower doses than used herein. Nevertheless, it is possible that the lower threshold for impact on birth weight in mammalian models could arise from maternal or placental effects, which are absent in the chicken embryo. This highlights how an advantage of the chicken embryo model, *in ovo* development, can present limitations in translation to human research. A key step in overcoming barriers to translation could be to quantify the carboxyhemoglobin of exposed embryos, allowing a more direct comparison of exposures between chicken embryos, rodent models and humans (Baker & Tumasonis, 1972).

The data presented here suggest that incubating eggs within a closed environment affects developmental stage, albeit not significantly, and that this effect is consistent between conditions. This delay should therefore have limited impact as a confounding factor. However, it must be considered in experiments examining effects and events at specific developmental stages. In this case, the delay could be overcome by increasing the incubation time or temperature (Table 1).

Hypoxia must be considered when incubating aerobic organisms in sealed environments. In this study we characterized the hypoxia arising from the longest period of closed box incubation. The observed levels remained above 17% O_2_. This is encouraging as the majority of studies identifying adverse negative effects of hypoxia in chick embryos use <17% O_2_ as their hypoxic condition (Lourens et al. 2007; Molenaar et al. 2010; Nangsuay et al. 2021). However, reduced body mass has also been described with incubation at 19% O_2_ from D18 to hatching (Wineland et al. 2006). While this exposure timing is not comparable to that of the present study, this nevertheless indicates that some manner of developmental delay may occur within a minimally hypoxic environment, potentially in line with the slightly delayed chick embryo development observed herein. Thus, establishing an acceptable range of hypoxia for novel experiments, based on the literature and experimental aims, might be prudent. We observe reduced hypoxia when incubating 2 rather than 3 eggs per box, offering a simple method for hypoxia reduction where experimentally necessary.

In conclusion, we present a novel method for the exposure of chicken embryos to gaseous and aerosol toxins. This method is non-invasive, affordable, accessible and technically straightforward in comparison to previously used methods. As with any protocol, there are important considerations for its use, particularly for experiments relating to hypoxia or specific developmental stages. While this method is validated for use with chicken embryos, it is suitable for exposure of smaller biological models, such as *Drosophila* or cultured mammalian cells. Using our protocol to continuously expose chicken embryos to up to 200ppm CO, we did not identify any significant effects of CO on embryo viability, weight or developmental stage. These findings are in agreement with previous research in chicken embryos, although a limited number of studies makes it difficult to draw definitive conclusions. It has long been understood that air pollutants pose a threat to human health, and epidemiological evidence suggests teratogenic effects in many cases. The method presented here provides a suitable platform for future work to develop our understanding of the teratogenic doses, mechanisms and effects of these air pollutants.

